# Characterisation of changes in the gut microbiota associated with eating disorders

**DOI:** 10.1101/2025.06.30.662422

**Authors:** Najate Achamrah, Jonathan Breton, Marie Galmiche, Thomas Demangeat, Muriel Quillard, Sébastien Grigioni, Vanessa Folope, Hélène Lelandais, Moïse Coëffier, Laure B. Bindels, Pierre Déchelotte, Marie-Pierre Tavolacci, David Ribet

**Author notes:** Corresponding author, INSERM UMR1073 – Université de Rouen, UFR Santé - 22 Boulevard Gambetta, 76183 ROUEN CEDEX. These authors contributed equally to this work (Najate Achamrah, Jonathan Breton).

## Abstract

**Background:** Eating disorders are serious pathologies that often begin in adolescence or young adulthood and persist for a significant period of time, with a strong negative impact on patients quality of life and mortality. The etiological origins of eating disorders are complex and involve both biological, psychological and societal factors. The gut microbiota was recently proposed as one of the potential factors involved in eating disorders. To gain a better understanding of the potential role of the gut microbiota in these diseases, we used 16S rRNA sequencing to compare the composition of the faecal microbiota of patients with all typical forms of eating disorders, i.e. anorexia nervosa, bulimia nervosa or binge-eating disorder, with that of healthy individuals.

**Results:** Our results demonstrate that each type of eating disorder is associated with a specific gut bacterial signature. We observed, for example, a decrease in the relative abundances of *Agathobacter* and *Romboutsia* genera and an increase in *Pseudomonas* in patients with anorexia, while patients with binge-eating disorder exhibit a decrease in the relative abundances of *Akkermansia* and *Intestinimonas* and an increase in *Streptococcus*, *Eggerthella* and *Proteus*. We also highlight a heterogeneity of gut microbiota composition in different subcategories of eating disorders, such as restricting versus binge-purge type anorexia or typical versus atypical binge-eating disorder. By focusing on the comorbidities reported by patients, we finally identified several bacterial taxa, such as *Acidaminococcus* and *Eggerthella*, whose level correlates with the occurrence of anxiety or depressive-like symptoms.

**Conclusions:** Together, our work demonstrates that eating disorders are associated with specific changes in gut microbiota composition and highlight the necessity to finely stratify patients to identify robust microbial signatures. In addition, we identified bacterial taxa correlating with comorbidities and decreased quality of life reported by patients. Our results now pave the way for determining the predictive value of the abundance of these taxa on the duration of the pathology or on the likelihood of relapse. They also constitute a valuable resource to further demonstrate the causal role of the gut microbiota in the onset or chronicisation of eating disorders.

## BACKGROUND

Eating disorders (EDs) are characterized by severe disturbances in eating behaviour and body weight. These disorders, which often begin in adolescence or young adulthood, are a major public health problem as they have a significant impact on patients’ quality of life and mortality [1]. Three typical EDs are defined according to the DSM-5 classification criteria: anorexia nervosa (AN), bulimia nervosa (BN) and binge-eating disorders (BED). Additional atypical or unspecified forms of eating disorders have also been described [2]. The lifetime prevalence of EDs has rapidly increased over the last twenty years, reaching around 8.4% in women and 2.2% in men [3]. Eating disorders are frequently associated with numerous psychiatric and somatic complications such as functional gastrointestinal disorders, anxiety and depression, which are shared among all types of EDs [4–6].

The molecular mechanisms that underlie EDs are still poorly understood. The etiology of these illnesses is multifactorial and involves both biological, psychological and societal factors. It has recently been proposed that the gut microbiota could be one of these factors. Indeed, the gut microbiota plays essential roles in digestion, energy extraction from food and weight regulation [7]. It may also regulate eating behaviour by modulating the secretion of intestinal appetite-regulating hormones or by sending signals to appetite-related neuronal circuitries in the central nervous system through the gut-brain axis [8, 9]. The gut microbiota finally impacts the hedonic and motivational aspects of food intake and regulates food reward responses [10].

Changes in the environment, diet or stress can modify the composition or metabolic functions of the gut microbiota, which can in turn affect the regulation of eating behaviour [8]. Several clinical studies explored whether patients with EDs exhibit intestinal dysbiosis. However, most of these studies were carried out on patients suffering from anorexia nervosa and very few focused on bulimia nervosa or binge-eating disorders [11–15]. In order to provide a detailed comparison of the gut microbiota composition in all typical EDs in parallel, we performed microbiome phylogenetic profiling on a cohort of patients suffering from anorexia nervosa, bulimia nervosa or binge eating disorders, as well as healthy individuals. Stratification of patients according to different clinical parameters and personal traits allowed us to identify bacterial taxa potentially involved in the onset or maintenance of EDs and their associated comorbidities.

## METHODS

### Study design

Patients over 18 of age, attending a first medical consultation at the Rouen University Hospital Nutrition Department, and with a clinician-diagnosed ED (according to the DSM-5 criteria) were eligible for inclusion and invited to participate to this study. Healthy volunteers over 18 years of age, with a Body Mass Index (BMI) between 18.5 and 24.9 kg/m^2^ and with no active or previous history of ED were eligible for inclusion and invited to participate to this study through the Rouen University Hospital Clinical Investigation Center (CIC-CRB 1404). Patients and healthy volunteers agreeing to participate provided a written informed consent and received a stool collection kit. Stools were collected at home and kept at 4°C before being transferred to the Rouen University Hospital Clinical Investigation Center within 24 hours, where they were immediately frozen at −80°C. Patients were grouped in three main categories of EDs: the “anorexia” group (including patients diagnosed with restricting type anorexia, binge/purge type anorexia, atypical anorexia or patients with restrictive dietary habits), the “bulimic” group (including patients diagnosed with typical or atypical bulimia nervosa) and the “binge-eating” group (including patients diagnosed with typical or atypical Binge-Eating Disorder and patients with intermittent compulsing eating behaviour or night eating syndrome) [16,17]. Body mass indexes were calculated using the weight and height measured on the day of inclusion.

### Questionnaires

Patients and healthy volunteers included in this study filled several questionnaires in order to collect data on both clinical features and personal traits or comorbidities. All questionnaires were translated in french.

*SCOFF questionnaire:* the SCOFF (Sick, Control, One stone, Fat, Food) questionnaire is classically used in primary care to screen for EDs. 2 positive answers or more (out of a total of 5 questions) predict the possible presence of an ED [18,19].

*HADS*: the HADS (Hospital Anxiety and Depression Scale) is based on a questionnaire containing two sets of 7 items, which evaluates individuals’ anxiety and depression states, respectively [20]. Subscores (HADS Anx and HADS Dep scores) were calculated to evaluate anxiety and depression independently. Individuals with subscores above 10 were considered as highly likely to have anxiety/depression-like syndromes [6].

*Rome III questionnaire*: this questionnaire aims to evaluate the occurrence of Irritable Bowel Syndrome (IBS). The diagnostic criteria for IBS are the occurrence of recurrent abdominal pain or discomfort (at least 3 days per month in the last 3 months, with symptom onset at least 6 months prior to diagnosis), associated with two or more of the following: improvement with defecation, onset associated with a change in frequency of stool, onset associated with a change in form (appearance) of stool [21].

*SF36 questionnaire*: The SF36 questionnaire is a generic quality of life questionnaire, widely used to obtain physical and mental health status [22]. This questionnaire is based on 36 items measuring eight quality of life dimensions : general health (SF36-GH), vitality (SF36-VT), bodily pain (SF36-BP), limitation of physical problems (SF36-RP), limitation of emotional problems (SF36-RE), mental health (SF36-MH), physical functioning (SF36-PF) and social functioning (SF36-SF). Subscores for each dimension were calculated, as well as two composite scores: the physical composite score (PCS) and the mental composite score (MCS), for which the PF, RP, BP and GH subscores and the VT, SF, RE and MH subscores were averaged, respectively [22]. Low SF36 scores are associated with a poor quality of life.

*EDI-2 questionnaire*: the EDI-2 (Eating Disorder Inventory-2) questionnaire is a standardized 91 items questionnaire initially designed for the assessment of anorexia nervosa and bulimia nervosa [23]. Based on the answers to this questionnaire, a global score and subscores focusing on drive for thinness (DT), bulimia (B), body dissatisfaction (BD), ineffectiveness (I), perfectionism (P), interpersonal distrust (ID), interoceptive awareness (IA), maturity fears (MF), asceticism (AS), impulse regulation (IR) and social insecurity (SI) were calculated. High EDI-2 scores are associated with a poor quality of life. This questionnaire has been filled by ED patients only.

*QUAVIAM questionnaire*: the QUAVIAM (Qualité de vie dans l’anorexie mentale) questionnaire is a 61 items-long french questionnaire developped initially to evaluate the quality of life of anorectic patients [24]. A global score is calculated for each patient. High QUAVIAM scores are associated with a poor quality of life. This questionnaire has been filled by ED patients only.

*BSQ questionnaire:* the BSQ (Body Shape Questionnaire) is a standardized 34 items questionnaire assessing individuals’ worries about weight and shape of the body [25,26]. A global BSQ score as well as 4 subscores were calculated from this questionnaire, that specifically focus on “avoidance and social shame of body exposure” (ABE), “bodily dissatisfaction with the lower parts of the body” (BDL), “use of laxatives and vomiting to reduce body dissatisfaction” (ULV) and “cognitions and inappropriate behaviours to control weight” (CIW) [25]. High BSQ scores are associated with significant bodily concerns. This questionnaire has been filled by ED patients only.

### Microbiota analysis

DNA from fecal samples was extracted as described previously [27]. The integrity of extracted DNA were assessed by visualizing their migration patterns on 1% agarose gels. Illumina sequencing of the 16S rRNA gene was done at the University of Minnesota Genomics Center. Briefly, the V5-V6 region of the 16S rRNA gene was PCR-enriched using the primer pair V5F_Nextera (TCGTCGGCAGCGTCAGATGTGTATAAGAGACAGRGGATTAGATACCC) and V6R_Nextera (GTCTCGTGGGCTCGGAGATGTGTATAAGAGACAGCGACRRCCATGCANCACCT) and then underwent a library tailing PCR as previously described [28]. The amplicons were purified, quantified and sequenced using an Illumina MiSeq to produce 2 x 300 bp sequencing products.

Initial quality-filtering of the reads was conducted with the Illumina Software, yielding an average of 76975 pass-filter paired reads per sample. 11 samples, yielding less than 2,000 reads, were removed from analysis. Raw sequences for the remaining samples can be found in the SRA database (Bioprojet: PRJNA1267983). Quality scores were visualized and reads were trimmed to 220 bp (R1) and 200 bp (R2) with the FASTX-Toolkit (http://hannonlab.cshl.edu/fastx_toolkit/). The reads were merged with the merge-Illumina-pairs application v1.4.2 [29]. The UPARSE pipeline implemented in USEARCH v11 was used to further process the sequences [30]. 7023 Amplicon sequencing variants (ASVs) were identified using UNOISE3 [31]. Taxonomic prediction was performed using the *nbc_tax* function, an implementation of the RDP Naive Bayesian Classifier algorithm [32] and the RDP database V16. Taxonomic identification of specific ASVs was further refined using the 16S rRNA gene database from EZbiocloud [33]. This led us in particular to reclassify the *Lachnospiracea_incertae_sedis* genus as *Agathobacter*. The phylotypes were computed as percent proportions based on the total number of sequences in each sample. Indexes of alpha diversity were computed using QIIME [34] on the rarefied ASV table. Rarefaction was performed using Mothur 1.32.1 [35] by randomly selecting a subset of 22072 reads for all samples except three (TCA1010: 16728; TCA1021: 13679; TCA1112: 10039).

Unrarefied taxa and ASV data were filtered to select for a minimum mean abundance in the whole cohort of 0.01% and a minimal prevalence of 10% in at least one group, leading to the selection of 937 ASVs. Permutational multivariate analysis of variance using distance matrices (ADONIS) was performed on centered log ratio (CLR)-transformed data, using the *adonis* function in the *vegan* v2.5-7 R package, to evaluate the contribution of a specific trait to the microbiome variance. A pseudo-count equal to half the minimal value found in the dataset was applied prior the CLR transformation [36]. The homogeneity of variances between groups was assessed using the *betadisper* function of the package *vegan* v2.5-7. Principal component analyses (PCA) were performed using the *pca* function of the mixOmics package [37]. Significantly impacted bacterial taxa were identified using a Mann-Whitney U-test. An alternative method to identify differential abundances, based on ALDEx2 [38], was used in parallel. For both methods, the obtained p-values were adjusted to control for the false discovery rate (FDR) for multiple testing according to the Benjamini and Hochberg procedure. q-values<0.1 were considered significant. Heat-maps were generated with the *heatmaply* R package, after normalization of the relative abundances in ED subtypes for each taxon. Correlograms were generated with the *corrplot* R package.

### Statistical analysis

Statistical analyses were performed using GraphPad Prism 8.0 and R. If normality was not respected in one group, Mann-Whitney test or Kruskal-Wallis test with Dunn’s correction were used to compare two groups or more than two groups, respectively. If normality was respected in all groups, two-tailed Student’s t-test or one-way ANOVA tests were used. Comparison of sex ratio and prevalences between two groups were performed using Fisher’s exact test. For all these tests, p-values<0.05 were considered statisticaly significant. Correlations between bacterial taxa or ASV and clinical parameters were performed by computing partial Spearman rank-based correlations, adjusted for potential covariates such as age, sex or BMI, using the *pcor* function in R (http://www.yilab.gatech.edu/pcor). FDR correction was applied. Bacterial taxa/ASV with correlations with a q-value<0.05 were considered statistically significant and included in heat-maps.

## RESULTS

### Clinical features of ED patients

181 patients from the EDILS (Eating Disorders Inventory and Longitudinal Survey) cohort [6,16,39], as well as 73 healthy individuals (HC), were included in this analysis. EDILS patients were grouped in 3 broad categories: (1) the “anorexia” group (AN; 35 patients, including 20 patients with restricting Anorexia, 6 patients with binge/purge type Anorexia, 1 patient with atypical Anorexia, and 8 patients with restrictive dietary habits); (2) the “bulimia” group (BN; 18 patients, including including 13 patients with typical Bulimia Nervosa and 5 patients with atypical Bulimia Nervosa) and (3) the “binge-eating” group (BED; 128 patients, including 74 patients with typical Binge-Eating Disorder, 31 patients with atypical Binge-Eating Disorder, 12 patients with intermittent compulsing eating behaviour and 11 patients diagnosed with Night Eating Syndrome) [17].

The population of ED patients was mainly composed of females (86%), which is consistent with the sex ratios classically observed for eating disorders (Table S1) [3]. Healthy individuals were selected for this study in order to have a similar sex ratio (*p*=0.35, Fishers’s exact test) and a similar mean age (*p*=0.23, Mann-Whitney test) compared to ED patients (Table S1). As expected, compared to healthy individuals, the mean Body Mass Index (BMI) is significantly lower in the “anorexia” group and significantly higher in the “binge-eating” group. No significant BMI differences is observed between the “bulimia” and the “healthy controls” groups (Table S1 and Fig. 1A).

**Figure 1:**
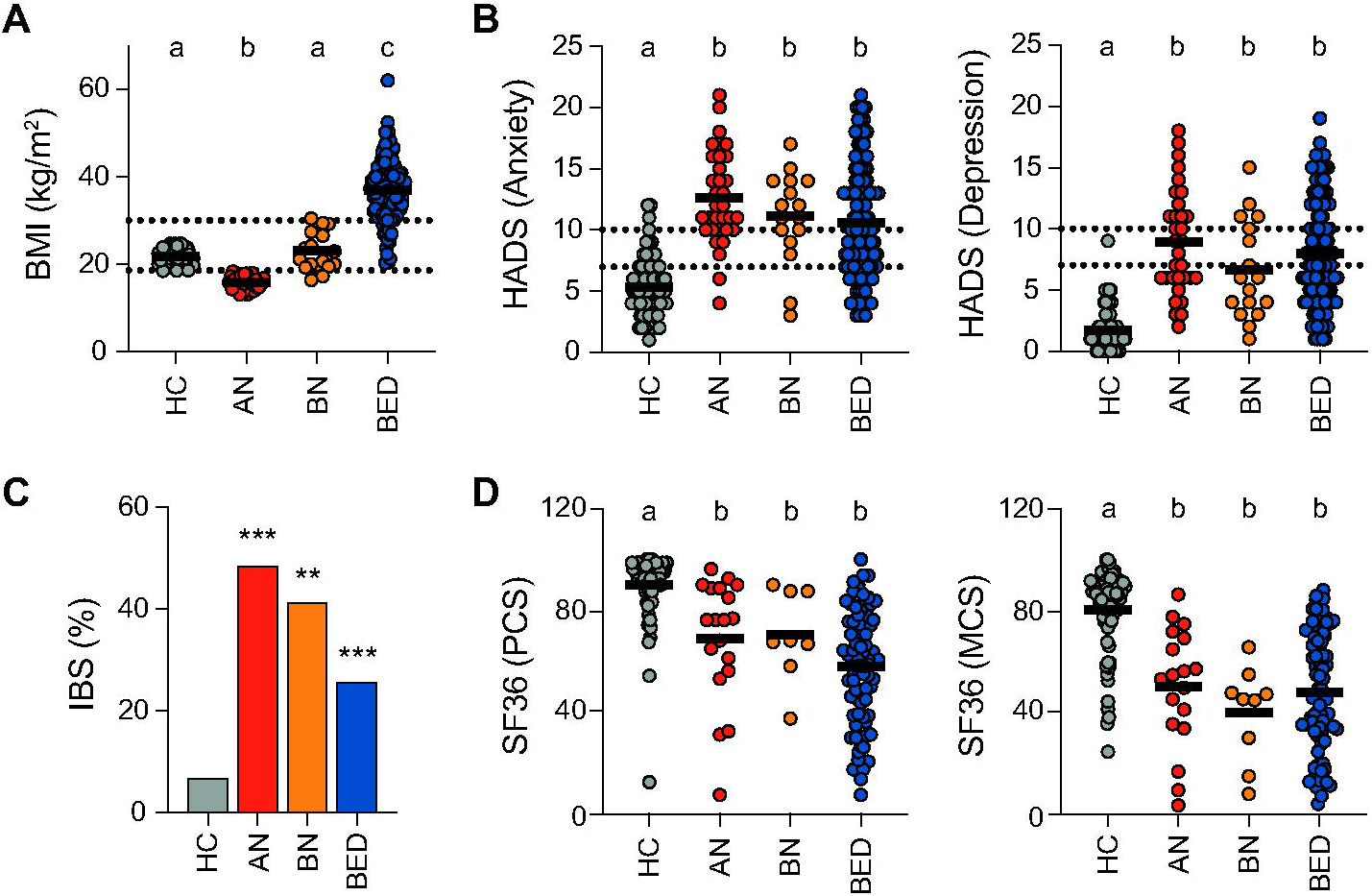
Clinical features of ED patients and healthy controls included in the study. A, Body Mass Index (BMI). BMI thresholds of 18.5 and 30 kg/m^2^, delineating under- and overweight individuals, respectively, are indicated by dashed lines. Individual values and means are represented (labeled means without a common letter differ; Kruskal-Wallis with Dunn’s correction). B, HADS scores for anxiety and depression. Score thresholds of 7 and 10, above which anxiety or depression syndromes are considered as likely or highly likely, respectively, are indicated by dashed lines. Individual values and means are represented (labeled means without a common letter differ; Kruskal-Wallis with Dunn’s correction). C, Percentage of individuals with IBS-like symptoms (**, p<0.01; ***, p<0.001; Fisher’s exact test). D, physical (PCS) and mental (MCS) composite scores calculated from the SF36 questionnaire. Individual values and means are represented (labeled means without a common letter differ; Kruskal-Wallis with Dunn’s correction). HC, healthy controls; AN, «anorexia» group; BN, «bulimia» group; BED, «binge-eating» group.

Several questionnaires were filled by ED patients and healthy individuals to get more insights into their personal traits and to detect potential comorbidities. As expected, ED patients show a SCOFF score significantly higher than healthy individuals (2.1±1.2 vs 0.1±0.2, *p*<0.001; Mann-Whitney test). Analysis of the HADS questionnaire revealed a significant increase in both the anxiety and depression scores of all ED groups compared to healthy individuals, with no significant differences between ED types (Fig. 1B). Anxiety (HADS anxiety score>10) was detected in 66%, 56% and 42% of patients within the “anorexia”, “bulimia” and “binge eating” groups, respectively, and in only 4% of healthy individuals. Depression (HADS depression score>10) was detected in 34%, 22% and 27% of patients within the “anorexia”, “bulimia” and “binge eating” groups, respectively, and not detected in any healthy individuals. This result confirms the high prevalence of anxiety and depression-like syndrome in patients with ED, which may worsen clinical symptoms [6,40–42]. Analysis of the Rome III questionnaire highlighted that the prevalence of functional gastrointestinal disorders is significantly increased in all ED groups compared to healthy individuals, as previously reported (Fig. 1C) [4,6]. Answers to the SF36 questionnaire, which focuses on patients’ quality of life, showed that all ED groups had a significant decrease in both the physical composite score (PCS) and the mental composite score (MCS) compared to healthy individuals, thereby highlighting a general decrease in the physical and mental health status of these patients, and thus in their quality of life (Fig. 1D) [22]. Of note, all the other SF36 subscores are decreased in at least one group of ED in comparison to healthy individuals (Fig. S1). We finally compared the answers of patients suffering from different type of EDs to the QUAVIAM, EDI-2 or Body Shape Questionnaires. We observed that the Body Shape Questionnaire (BSQ) global score is significantly increased in the “bulimia” or “binge-eating” groups in comparison to the “anorexia” group, thus indicating more severe bodily concerns in these two first categories of ED patients (Table S2). This result is consistent with previous studies showing that the BSQ global score is positively correlated with BMI [25]. No significant differences were observed in the global QUAVIAM and EDI-2 scores between ED types (Table S2).

### Characterization of gut microbiota changes associated with each type of eating disorder

Fecal samples from healthy individuals and patients with EDs were collected in order to compare the composition of the gut microbiota between these individuals. DNAs were extracted from these samples and microbiome phylogenetic profiling was performed using 16S rRNA gene amplicon sequencing.

We first estimated the effect size of various parameters on the composition of the gut microbiota from our studied population using PERMANOVA. We observed that the type of ED has the largest explanatory power on gut microbiota composition among all tested factors (explaining 2.6% of ASV abundance variations) (Fig. 2). We also found a significant effect size for BMI (1.6%). The effect size of these factors remains limited, as additional unknown factors or stochastic effects are probably also contributing to the composition of the gut microbiota of these individuals. We also demonstrate that the use of medication has a significant impact on microbiota composition, and more particularly the use of antidepressants, anxiolytics and vitamins during the past 12 months, or the current use of nutritional supplements (Fig. 2). Previous medical history of malabsorptive syndrome, depression or anxiety also have a significant effect size (Fig. 2).

**Figure 2:**
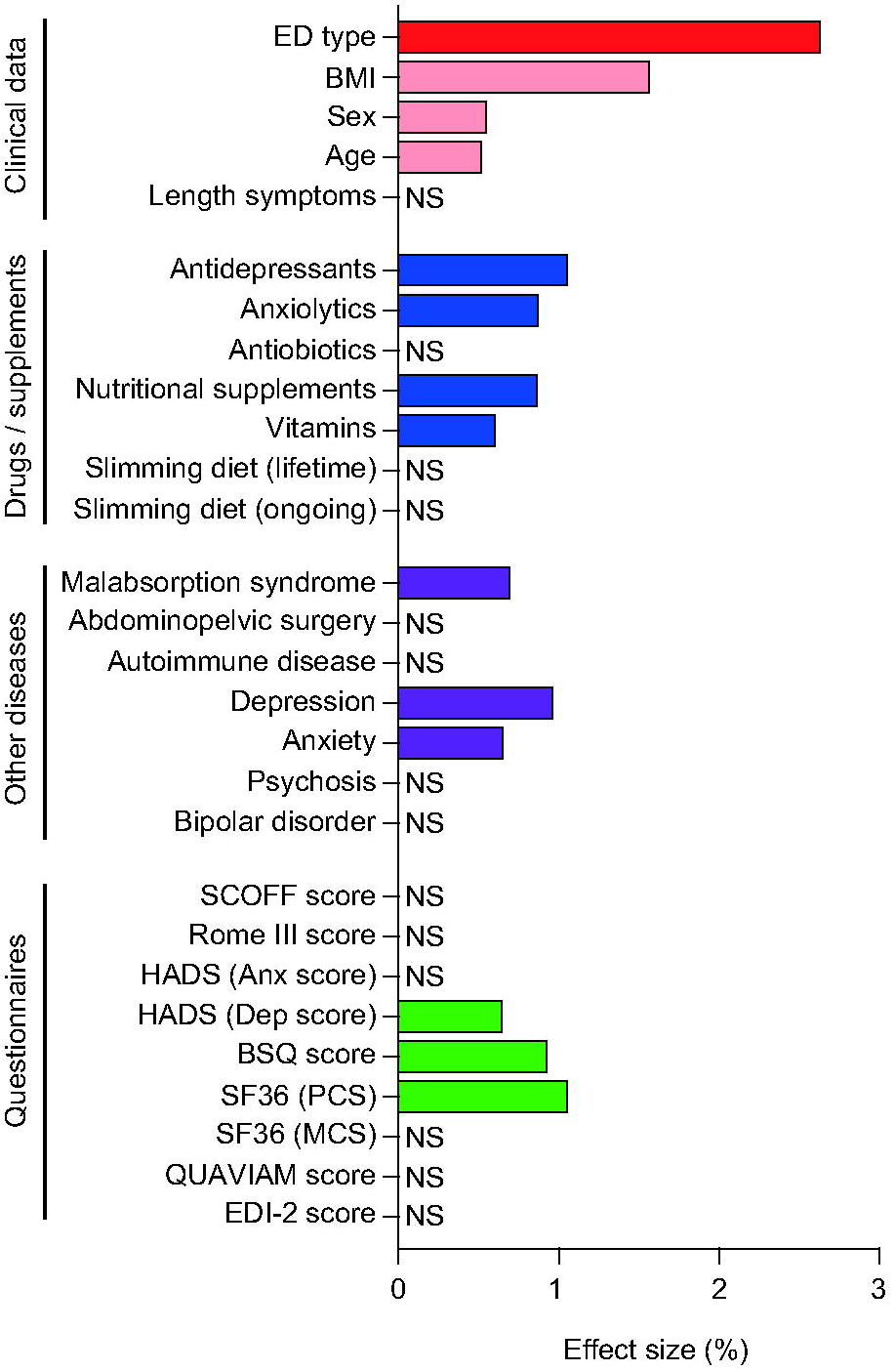
Effect size of various clinical parameters on the overall gut microbiota composition. Effect size was determined by ADONIS analyses on ASV data (NS, not significant).

In order to compare the gut microbiota composition between the different types of eating disorders, we first calculated alpha-diversity indexes to estimate within samples richness and evenness. A significant decrease in the number of observed species was observed for patients in the “binge-eating” group compared to healthy individuals, but not for other EDs (Fig. S2). No significant differences were observed for Shannon, Simpson, Heip evenness and Simpson evenness indexes between ED patients and healthy individuals (Fig. S2). These results suggest that the gut microbiota’s richness and evenness is not altered in patients suffering from anorexia or bulimia, and that gut microbiota’s richness is decreased in patients suffering from binge-eating disorder.

We then compared the composition of the gut microbiota between ED patients and healthy individuals at the ASV level (Fig. 3A). PERMANOVA demonstrates that the overall gut microbiota composition is significiantly dissimilar according to the type of ED (R^2^=2.6%, p=0.001). Of note, a significant difference in dispertion was observed between healthy individuals and the “anorexia” group (p=0.017, betadisper), but not between healthy individuals and the “bulimia” or “binge-eating” groups. Dissimilarities in gut microbiota composition were similarly observed when performing PERMANOVA at different taxonomic levels, including families and genera.

**Figure 3:**
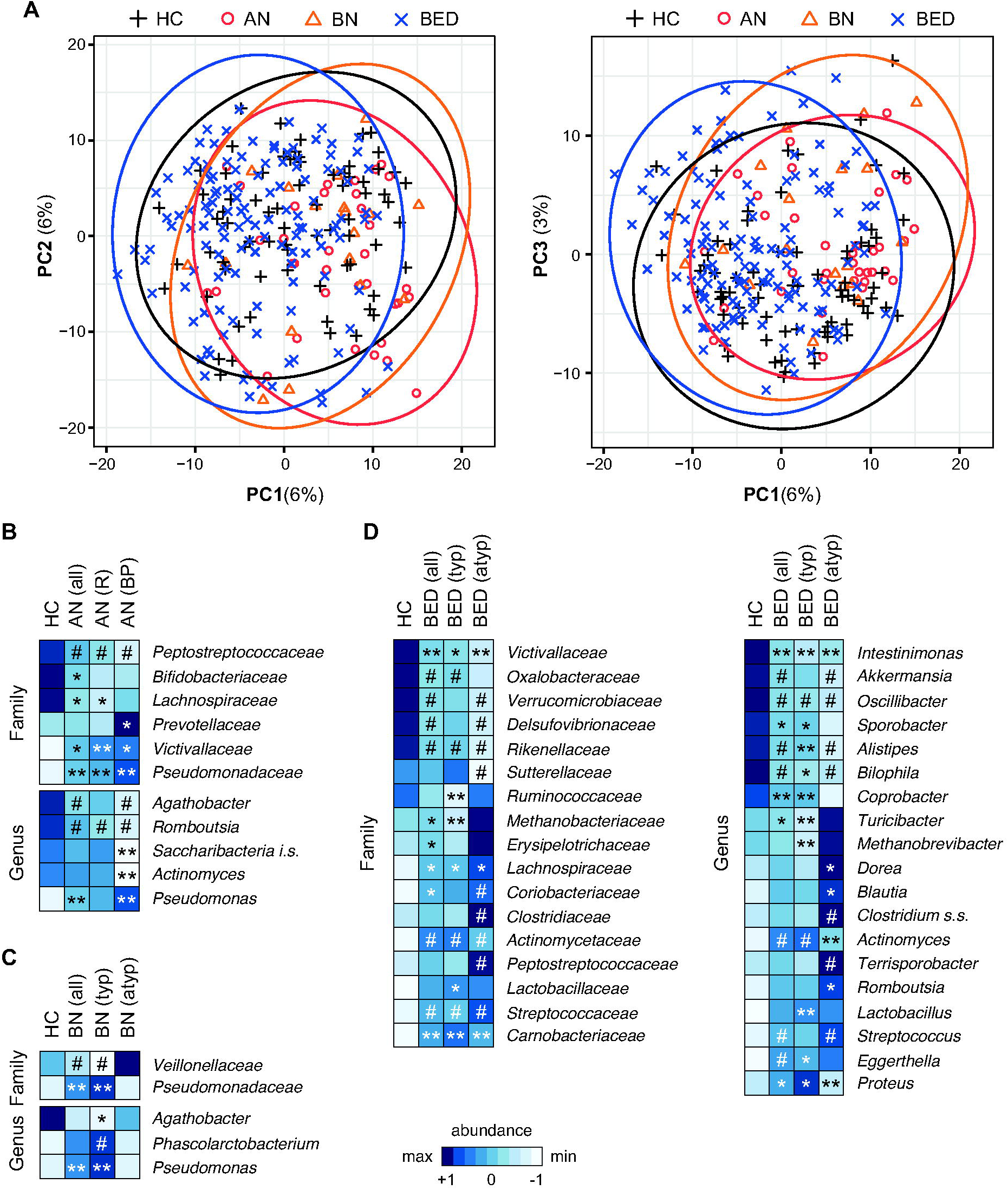
Characterization of the gut dysbiosis associated with ED. A, Principal Component Analyses of the gut microbiota composition in ED patients and healthy individuals. (B-D) Heat-map of bacterial families and genera showing altered relative abundances between ED patients and healthy individuals(*, q<0.1; **, q<0.05; Mann-Whitney;#, q<0.1 for both Mann-Whitney and Aldex2). HC, Healthy controls; AN, «anorexia» group; BN, «bulimia» group; BED, «binge-eating» group. AN(R), Anorexia-restricting type; AN(BP), Anorexia-Binge/purging type; typ, typical; atyp, atypical. Colors are assigned based on transformed abundances after mean normalization.

To gain further insights into the changes in microbiota composition in ED patients, we compared the relative abundance of 937 ASVs in ED patients and healthy individuals. We observed that 56 ASVs in AN patients, 2 ASVs in BN patients and 110 ASVs in BED patients showed significant altered relative abundances compared to healthy individuals (q<0.1), which confirms the occurrence of a gut bacterial signature specific to each type of ED (Table S3). The limited number of ASVs with altered abundances in BN patients is probably due to the smaller number of patients included in this group. In AN and BED patients, most of the identified ASVs exhibit decreased relative abundances compared to healthy individuals. By focusing only on the most abundant ASVs (with a relative abundance >0.5% in healthy controls), we identified 3 ASVs assigned to the *Agathobacter*, *Bifidobacterium* and *Gallintestinimicrobium* genera, that are significantly decreased in AN patients and 8 ASVs assigned to the *Akkermansia*, *Barnesiella, Dysosmobacter*, *Vescimonas* and *Waltera* genera and unclassified *Christensenellaceae* and *Oscillospiraceae*, that are significantly decreased in BED patients (Fig. S3). Of note, we carried out the same analyses on women only and obtained similar results (Table S3).

To complete these results, we performed similar analyses on higher taxonomical levels. We identified in total 55 bacterial taxa with increased or decreased relative abundances between ED patients and healthy individuals (Fig. 3B-D, Fig. 4 and Table S4). More particularly, we observed that patients in the “anorexia” group exhibit an increase in *Pseudomonas* and a decrease in *Romboutsia* and *Agathobacter* genera. Of note, patients from the “anorexia” group also exhibit an increase in the *Euryarchaeota* phylum, which comprises the methanogenic archaeon *Methanobrevibacter smithii* (Fig. 4 and Table S4). Patients in the “bulimia” group exhibit an increase in *Pseudomonas*, similarly to patients with AN. Finally, patients in the “binge-eating” group exhibit a significant decrease in several genera including *Akkermansia*, *Intestinimonas, Oscillibacter* and *Sporobacter* (fold-change versus healthy controls <0.67) and an increase in *Actinomyces*, *Eggerthella, Proteus* and *Streptococcus* (fold-change versus healthy controls >1.5) (Fig. 3B, Fig. 4 and Table S4). In addition to differences in relative abundances, we also noticed altered prevalence of specific bacterial taxa in ED patients. This is the case for example for the *Pseudomonas* genus, which has an increased prevalence in the “anorexia” and “bulimia” groups and the *Proteus* genus, which has an increased prevalence in the “binge eating” group (Fig. 4). Of note, the same analysis performed on women only resulted in similar results (Table S4).

**Figure 4:**
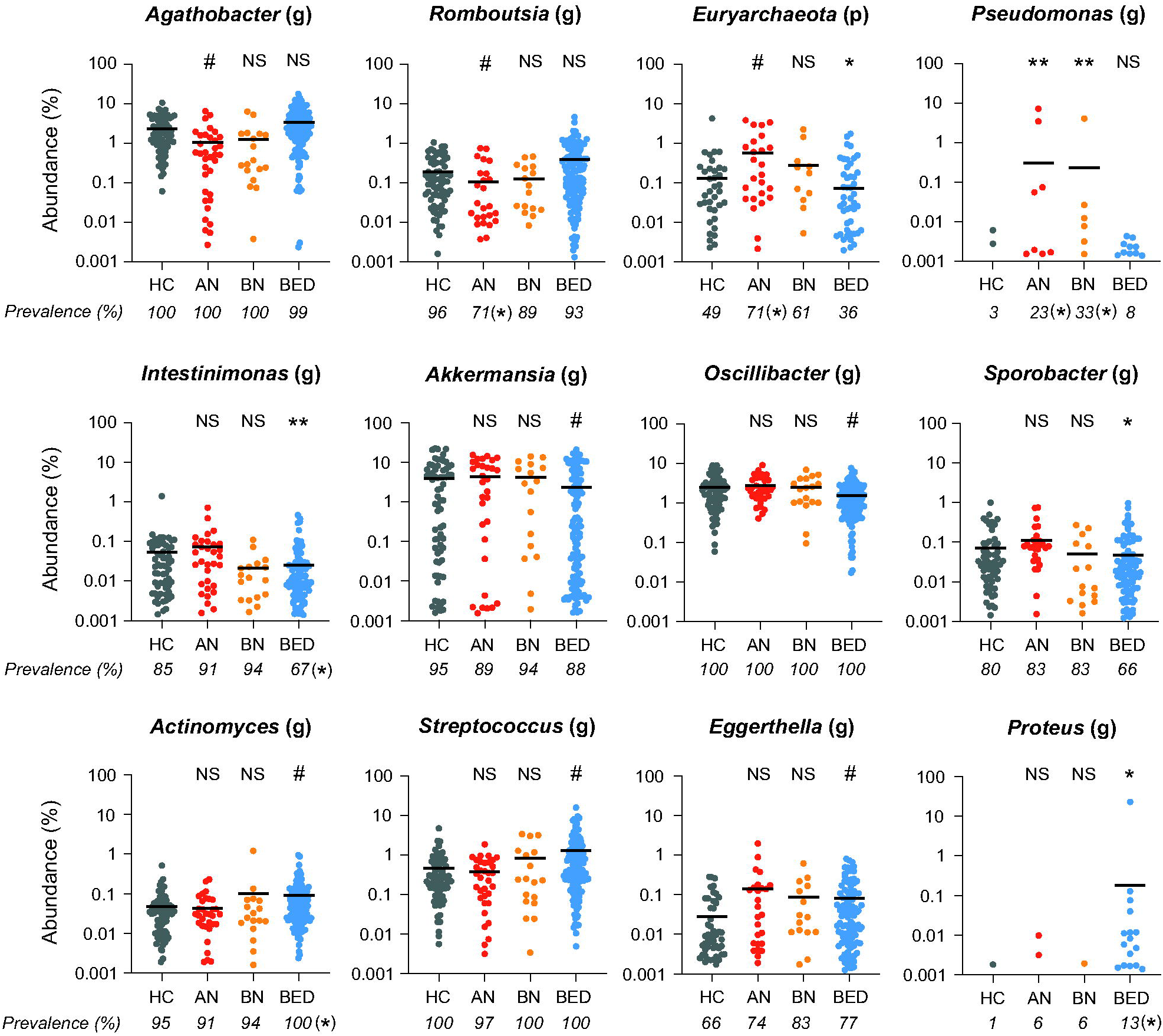
Relative abundances of bacterial taxa altered in ED patients. Examples of bacterial taxa showing significantly altered abundances in ED patients (*, q<0.1; **, q<0.05; Mann-Whitney; #, q<0.1; Aldex2; comparison versus HC). Prevalences in each group are indicated below graphs (*, p<0.05; Fisher’s exact test vs HC). HC, Healthy controls; AN, «anorexia» group; BN, «bulimia» group; BED, «binge-eating» group; (g) genus.

Together, these results demonstrate that each type of ED is associated with a specific gut bacterial signature, since the ASV/genera showing altered relative abundances are not overlapping between the groups.

Since the “anorexia”, “bulimia” and “binge-eating” groups used here encompass various phenotypes, we compared the relative abundances of bacterial taxa in these different subcategories (Fig. 3 and 5). Interestingly, we identified several bacterial taxa showing differential abundances between ED subcategories. For example, the *Actinomyces* genus is significantly decreased in patients with the binge-purge type anorexia but not in patients with restricting anorexia. Similarly, the *Romboutsia* genus is significantly increased in patients with atypical BED but not in patients with typical BED (Fig. 3 and 5). Of note, no significant differences were observed in the relative abundances of *Firmicutes* (newly reassigned as *Bacillota*) and *Bacteroidetes* (newly reassigned as *Bacteroidota*) phyla among the different ED subcategories (Fig. 5). Together, these results suggest that gut microbiota composition is heterogenous between the different ED subtypes.

**Figure 5:**
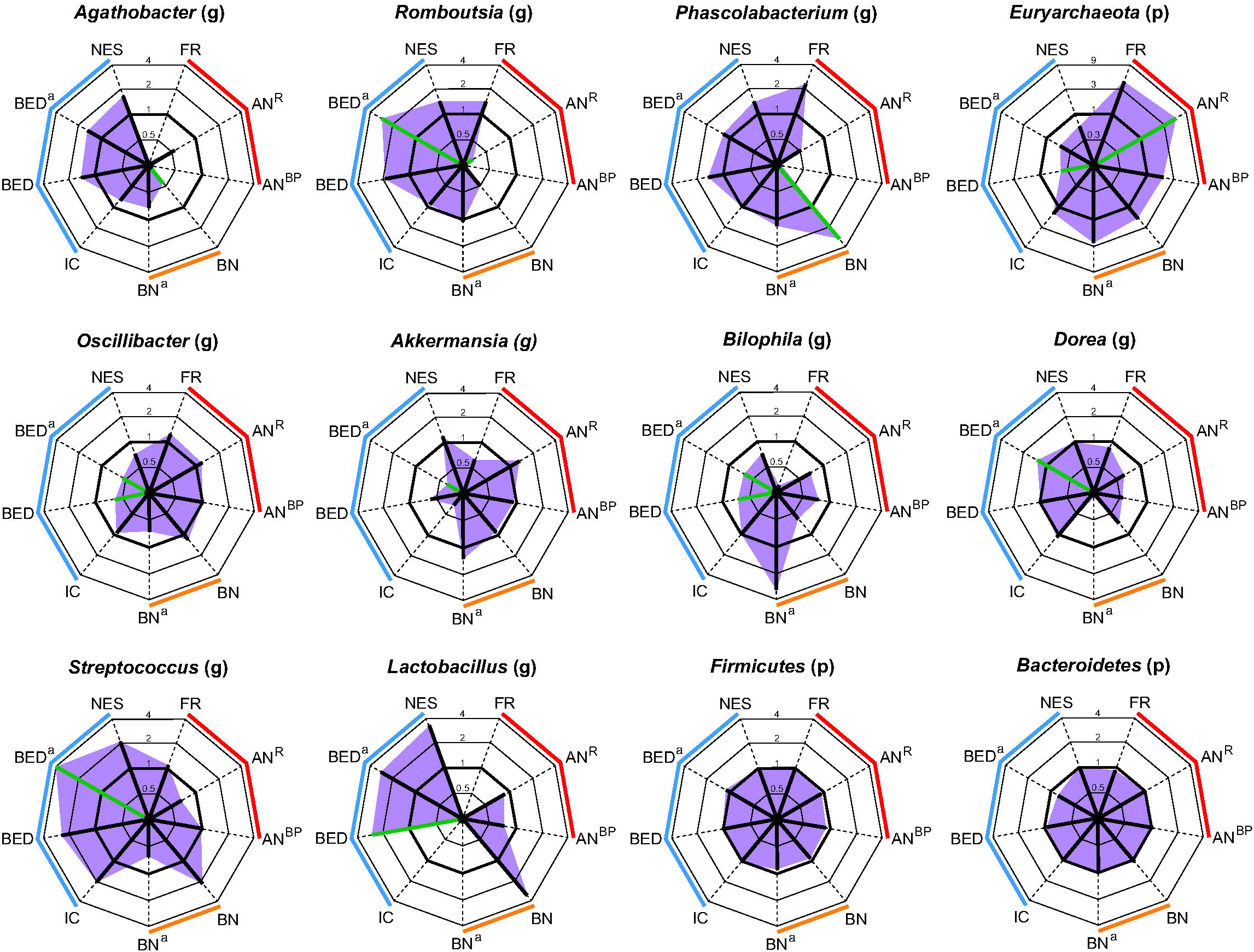
Relative abundances of bacterial taxa in specific ED categories. Radar charts of bacterial taxa in different categories of ED. Red, «anorexia» group; orange, «bulimia» group; blue, «binge-eating» group. FR, food restriction; ANR, Anorexia-restricting type; AN^8^P, Anorexia-binge/purging type; BN, typical Bulimia; BNa, atypical Bulimia; IC, intermittent compulsivity; BED, typical Binge-Eating Disorder; BEDa, atypical Binge-Eating Disorder; NES, Night Eating Syndrome. (p), phylum; (g), genus. Green lines correspond to significantly altered levels (q<0.1; Mann-Whitney; comparison versus healthy controls).

### Modification of the gut microbiota composition associated with ED comorbidities

We assessed whether comorbidities frequently observed in ED patients, *i.e.* functional gastrointestinal disorders (IBS-like symptoms), anxiety and depression, were correlated with a shift in gut microbiota composition. For this, we either compared the microbiota composition in patients reporting or not comorbidities, or performed partial Spearman ranked-based correlations between the relative abundances of bacterial taxa and scores from the HADS questionnaire. We performed correlations either on the global pool of healthy individuals and ED patients or in the “anorexia” and “binge-eating” groups separetely (the “bulimic” group was not analyzed since it includes less than 20 patients). Corrections for BMI, sex and age were performed. Similar analyses were run on women only.

We first compared the microbiota composition between patients reporting or not functional gastrointestinal disorders, in each type of eating disorder. ADONIS analysis did not identify dissimilarities at the genus or ASV levels between these groups. By looking at each taxon and ASV individually, in each type of ED, we identified one ASV, corresponding to a bacterium from the *Bacteroides* genus, which is specifically enriched and has an increased prevalence in patients with functional gastrointestinal disorders in the “binge-eating” group (Fig. 6A). In contrast, no ASV nor taxa show altered levels in patients reporting IBS-like symptoms in the “anorexia” and “bulimia” groups. To complete this analysis, we then stratified patients depending on the abundance of specific bacterial taxa. We observed that in the “anorexia” group, patients with high level of *Akkermansia* (abundance>0.5%) are more likely to report IBS-like symptoms than patients with low level of *Akkermansia* (p<0.05; Fisher’s exact test). This difference is specific to the “anorexia” group and not observed for healthy individuals or patients from the “bulimia” or the “binge eating” groups (Fig. 6B). No other stratification showed differences in the prevalence of IBS-like symptoms. Taken together, these results suggest that, within a given ED, patients with IBS-like symptoms exhibit only slight modifications in gut microbiota composition compared to patients without IBS-like symptoms.

**Figure 6:**
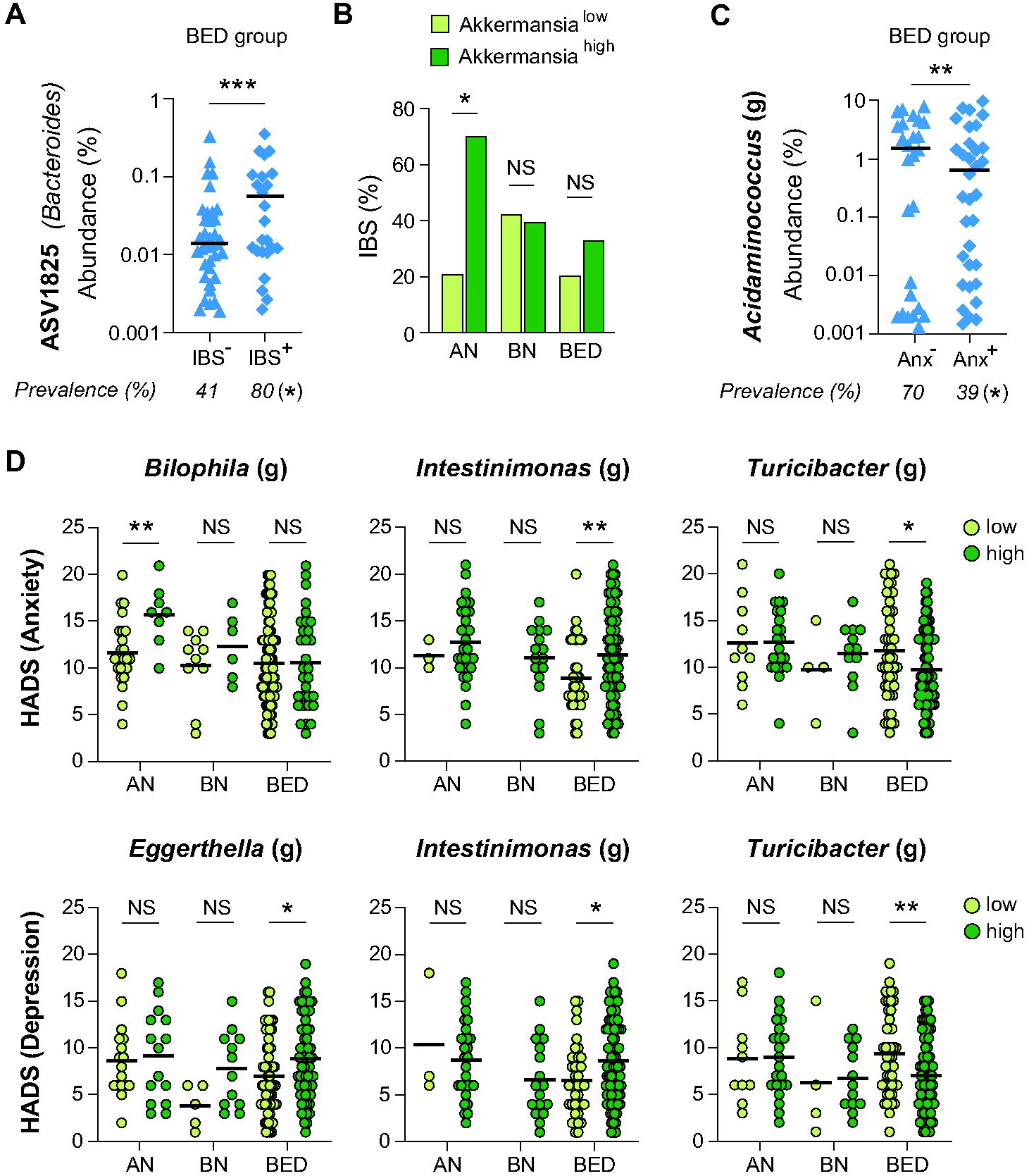
Bacterial taxa altered in ED patients with comorbidities. Examples of ASV (A) and bacterial taxa (C) showing significantly altered abundances in ED patients with comorbidities (**, p<0.01; ***, p<0.001; Mann-Whitney test). Prevalences in each group are indicated below graphs(*, p<0.05; Fisher’s exact). B, Percentages of individuals with IBS-like syndrom in patients with low or high levels of *Akkermansia* (*, p<0.05; NS, not significant; Fisher’s exact test). D, HADS anxiety and depression scores in patients with low or high levels of specific bacterial genera(*, p<0.05; **, p<0.01; NS, not significant; Mann-Whitney test). AN, «anorexia» group; BN, «bulimia» group; BED, «binge-eating» group. (g), genus.

In parallel to this study on functional gastrointestinal disorders, we focused on the differences in gut microbiota composition between patients reporting or not anxiety-like disorders. Of note, very few patients from the “anorexia” and “bulimia” groups are free from anxiety-like disorders (2/33 and 2/16 patients reported an HADS Anx score <=7 in the “anorexia” and “bulimia” groups, respectively). Thus, we focused on the “binge-eating” group (for which 37 patients reported an HADS Anx score <=7 and 54 patients a score >10) to compare microbiota composition in patients with or without anxiety-like disorders. PERMANOVA revealed dissimilarities at the genus level between the gut microbiota composition in patients from the “binge-eating” group with or without anxiety-like disorders (R^2^=0.017, p=0.024), as well as a difference in dispertion (p=0.04, betadisper). Interestingly, univariate analyses identified one bacterial taxon, the genus *Acidaminococcus*, with significantly decreased abundance and prevalence in binge-eating patients reporting anxiety-like symptoms compared to patients without anxiety (Fig. 6C). We further identified that the level of *Acidaminococcus* was negatively correlated with the HADS Anx score in our cohort and, more specifically, in the “binge-eating” group (rho = −0.23 and −0.34, respectively; q<0.05) (Fig. 7). We then performed analyses on patients stratified on the abundance of specific taxa. We observed that in the “anorexia” group, patients with high level of the *Bilophila* genus (abundance>0.3%) have a significantly higher HADS Anx score (Fig. 6D). In the “binge-eating” group, patients in whom bacteria from the *Intestinimonas* genus were detected exhibit higher HADS Anx scores, whereas patients with *Turicibacter* bacteria exhibit lower HADS Anx scores (Fig. 6D). These associations are not observed for other types of EDs. Together, these results show that specific bacteria, such as *Acidaminococcus*, correlates with the anxiety level in patients with EDs and that these associations are specific to the type of ED.

**Figure 7:**
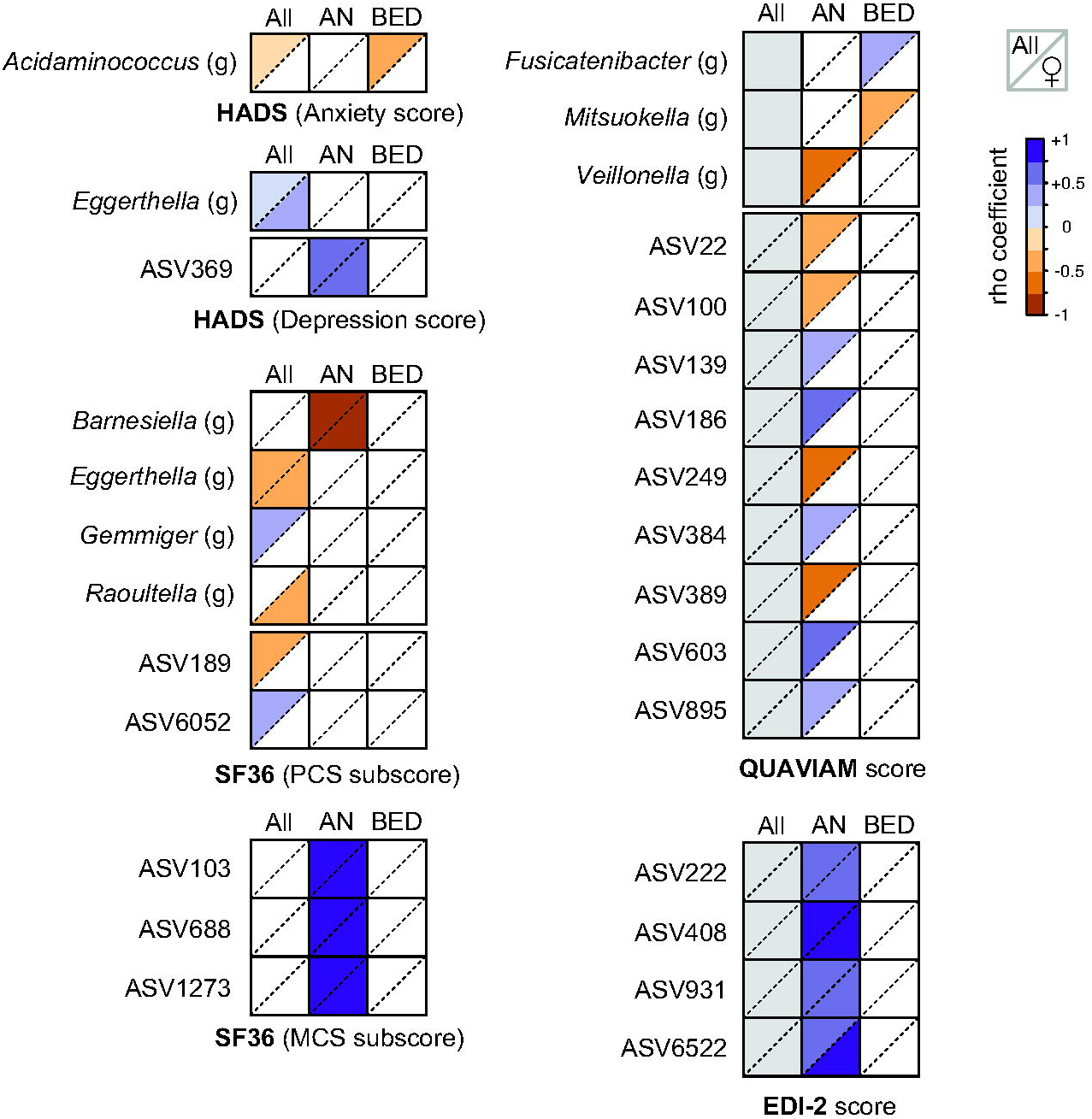
Correlation of bacterial taxa and ASV abundances with clinical scores. Partial Spearman ranked-based correlations between the relative abundances of bacterial taxa and ASV and scores from the HADS, SF36, QUAVIAM and EDl2 questionnaires. Only significant correlations are indicated (q<0.05). Correlations were performed on all individuals, including healthy controls (All), or in the «anorexia» (AN) and «binge-eating» (BED) groups separetely. Correlations were performed on both males and females (top left triangles) or on females only (bottom right triangles). Colors correspond to Spearman rho coefficients. (g), genus. No significant correlations were observed between bacterial taxa/ASV and BSQ scores.

We finally focused on depression-like symptoms. We performed PERMANOVA to decipher whether reporting of depression-like symptoms is associated with differences in gut microbiota composition within each type of ED. We did not identify any significant differences in the global microbiota composition, nor in taxa or ASV levels between patients exhibiting or not depression-like symptoms. Correlation analyses performed on the global cohort identified that the levels of *Eggerthella* positively correlates with the HADS Dep score (rho = +0.22; q<0.05) (Fig. 7). Stratification of patients based on the abundance of specific taxa confirmed that patients from the “binge-eating” group exhibiting high levels of *Eggerthella* (abundance>0.2%) have higher HADS Dep scores. Similar results were obtained for patients in whom bacteria from the *Intestinimonas* genus were detected, whereas patients with *Turicibacter* exhibit lower HADS Dep scores (Fig. 6E). Together, these results show that some specific bacteria are associated with high or low level of depression in patients with EDs.

Of note, we also performed correlations between bacterial abundances and BMI. We identified several genera and ASV which were correlated with BMI when we performed our analysis on the global cohort (after correction for sex and age; Table S5). However, we did not identify taxa or ASV correlating with BMI when we focused only on the “anorexia” or “binge-eating” group, except for ASV854 (corresponding to a bacterium from the *Lachnospiraceae* family), which relative abundance positively correlates with BMI in the “anorexia” group (Table S5).

### Correlations between bacterial genera and ASV abundances and patients’ quality of life

To complete these analyses on ED comorbidities, we looked for potential correlations between the levels of bacterial genera/ASV and the scores calculated from the QUAVIAM, EDI-2, BSQ or SF36 questionnaires. For the SF36 questionnaire, we performed correlations either on the global cohort or on the “anorexia” and “binge-eating” groups, separetely. For the QUAVIAM, EDI-2 and BSQ questionnaire, which were filled only by ED patients, we performed correlations only on the “anorexic” and “binge-eating” groups.

For the SF36 questionnaire, we observed that the abundance of the bacterial genus *Gemmiger* is positively correlated with the SF36 physical composite score (PCS), whereas several bacterial genera are negatively correlated with this score (q<0.05). These bacteria include *Eggerthella* and *Raoultella* (for the global cohort) and *Barnesiella* (specifically in the “anorexia” group) (Fig. 7). These last three genera are thus associated with a decreased quality of life concerning one or several of the following items: physical functioning, bodily pain, limitation of physical problems and general health. By performing similar correlation analyses on ASV levels, we identified 2 ASV which correlates with SF36 PCS score, and 3 ASVs correlating with SF36 mental composite score (MCS) (Fig. 7 and Suppl. Table S6). Concerning the QUAVIAM questionnaires, which evaluate the quality of life of ED patients, we also identified several genera whose abundance are negatively correlated with QUAVIAM scores. This is the case of *Veillonella*, in the “anorexia” group, and *Mitsuokella,* in the “binge-eating” group (Fig. 7). In contrast, we identified that the genus *Fusicatenibacter* is positively correlated with QUAVIAM scores in the “binge-eating” group. Knowing that a high QUAVIAM score is associated with poor quality of life, this indicates that the abundance of *Veillonella* correlates with a higher quality of life reported by AN patients. 9 ASVs were further identified has positively or negatively correlated with QUAVIAM scores in the “anorexia” group specifically. Concerning the EDI-2 questionnaires, 4 ASVs were identified as positively correlated with EDI-2 scores. This indicates that a higher abundance of these ASVs correlates with a poorer reported quality of life in patients. No significant correlations were observed between bacterial taxa/ASV abundances and BSQ scores.

Together, these results highlight that specific bacterial taxa or ASVs are specifically correlated with patient’s quality of life.

## DISCUSSION

Our study provides a detailed characterization of the gut dysbiosis occuring in eating disorders. A major strength of our study is the inclusion of patients suffering from all typical forms of EDs, thereby allowing the comparison of their gut microbiota with a unique methodological approach. In addition, patients from the EDILS cohort were included during their first medical consultation at the Rouen University Hospital Nutrition Department, which limits the potential bias related to previous therapeutic intervention.

Thanks to this study design, we demonstrate that each type of ED is associated with a specific gut microbial composition. Accordingly, we observed that the type of ED has the largest explanatory power on gut microbiota composition among all tested factors in our cohort. We identified as well an heterogeneity in the gut bacterial signatures between different subcategories of AN (binge-purge type or restricting), BN (typical or atypical) and BED (typical or atypical). These results suggest that a fine stratification of ED patients is essential to identify robust microbial signatures associated with these diseases. Lack of such stratification may contribute to the inconsistency in gut dysbiosis observed in the literature to date, particularly in the case of AN [11–15].

Gut microbiota analysis of the “anorexia” group from our cohort revealed an alteration of several bacterial taxa and ASV encompassing a decrease in *Agathobacter* and *Romboutsia,* as well as an increase in *Pseudomonas.* The observed decrease in the relative abundance of *Agathobacter* and *Romboutsia* in our “anorexia” group have similarly been reported in previous clinical studies [43–47]. The decrease in the abundance of *Agathobacter,* which are butyrate-producing bacteria, may participate to the decrease in butyrate levels observed in AN patients [48,49]. Butyrate is a short-chain fatty acid (SCFA) playing essential roles in the regulation of the intestinal barrier as well as in mood or behaviour. It has been proposed that the decrease in butyrate levels in AN patients could also be due to a decrease in *Roseburia* species [12,46,50]. We observed in our “anorexia” group a decrease in *Roseburia* abundance but it did not reach significance (x0.52 vs HC, p=0.02, q=0.13). In contrast to this decrease in butyrate-producing bacteria, we observed an increase in *Pseudomonas* abundance in our “anorexia” group, which is usually considered as a pathobiont. This overgrowth of pathobionts may exacerbate the impact of the loss of beneficial bacteria and participate to the alteration of gut functions in patients, such as the alteration in the intestinal barrier observed in severely malnourished AN patients [51]. Finally, an increase in *Methanobrevibacter* levels was frequently reported in clinical studies on AN [12,50]. We also observed in our “anorexia” group an increase in *Methanobrevibacter* but it did not reach significance (x2.91 vs HC, p=0.03, q=0.20). Of note, we observed a significant increase in the phylum *Euryarchaeota*, which includes the *Methanobrevibacter* genus, in patients with anorexia and more particularly with restricting AN. Besides this modification in bacterial taxa or ASV abundances, we also noted an increased dispersion in this group, suggesting a higher inter-individual variability in the microbiota composition of patients with AN, but did not observe any significant changes in alpha-diversity indexes in patients with anorexia compared with healthy individuals.

In the case of Bulimia Nervosa, we identified only few bacterial taxa and ASVs with altered levels between patients and healthy individuals. This low number of altered taxa/ASV may be due to the limited number of patients included in our analyses (18 BN patients in total). No significant changes in alpha diversity indexes was observed in patients with BN. We identified a decrease in *Agathobacter* and an increase in *Pseudomonas* relative abundances, specifically in patients with typical BN. Interestingly, these two bacterial genera are similarly affected in patients with binge-purge type AN from our cohort, but not with restricting AN. Alteration of these bacterial taxa thus correlate with the binge/purge conduct of these patients from two different ED subcategories. Three other studies addressed gut microbiota composition in patients with BN so far [14,15,52]. The grouping of patients in these studies is however highly heterogenous which makes it impossible to determine the characteristics of the core dysbiosis in BN patients. Further studies in larger populations of well-phenotyped patients are thus needed to obtain a more coherent characterization of the microbial signatures associated with BN.

In the case of patients with BED, we observed a global decrease in gut microbiota richness and a specific decrease in *Akkermansia*, *Intestinimonas, Oscillibacter* and an increase in *Actinomyces*, *Eggerthella, Proteus* and *Streptococcus* relative abundances. One previous study compared the gut microbiota composition in fecal samples between patients with both obesity and BED and patients with obesity without BED [13]. Interestingly, this study also identified a decrease in the abundances of *Akkermansia* and *Intestinimonas* in BED patients. *Akkermansia* is a well-established SCFA producer and a decrease in its abundance is usually linked with altered metabolic profile. Indeed, this bacterium was shown to improve gut-barrier function, reduce low-grade inflammation, improve sensitivity to insulin and decrease adiposity [53–55]. *Intestinimonas* is also a SCFA-producer and its decrease may be potentially harmful [56]. Accordingly, in a model of mouse with diet-induced obesity, oral supplementation with *Intestinimonas butyriciproducens* counteracts body weight gain, hyperglycemia and adiposity [57]. Concerning the *Proteus* genus, an increase of its prevalence was observed in colonic tissues of patients with Inflammatory Bowel Disease and bacteria from this genus were shown to induce inflammation in cells and animal model of colitis [58]. These bacteria may thus act as pathobionts and promote low-grade intestinal inflammation associated with BED. Several bacterial taxa were differentially altered in typical and atypical BED. We observed in particular that several genera such as *Dorea*, *Blautia*, *Terrisporobacter*, *Romboutsia* or *Streptococcus* showed increased relative abundances in atypical BED but not in typical BED. Again this results highlight the heterogeneity of the gut microbial signatures in different ED subtypes and the necessity to finely phenotype ED patients in order to delineate robust signatures associated with these diseases.

Of note, we did not compare gut microbiota composition in our BED group with that of a group of obese patients without BED. We cannot thus rule out that some of the bacterial taxa identified as deregulated in patients with BED are not specifically linked to this ED but are rather associated with increased BMI in these patients (36.9 ± 6.8 kg/m^2^).

We also focused in our study on gut microbiota alteration associated with ED-related comorbidities. Thanks to the different questionnaires included in our study, we confirmed the high prevalence of IBS-like symptoms in all types of ED, the significant increase in anxiety and depression-like symptoms and the global decrease in quality of life reported by patients [4,6,22,40,41]. We did not identify strong microbial signatures associated with IBS-like symptoms in the different ED categories. Unlike anxiety or depression which were assessed using a quantitative score, we only have a binary information about the presence of IBS-like symptoms in our cohort. The grouping of patients with potentially heterogenous severity or symptoms of IBS may have impaired our ability to detect specific bacterial signatures. Surprisingly, we noted that patients with high relative abundance of *Akkermansia* (>0.5%) were more likely to reports functional gastrointestinal disorders, specifically in AN patients. Potential associations between *Akkermansia* and IBS-related symptoms have already been reported in the literature [59–61]. However, care must be taken when interpreting these results on *Akkermansia*, which are merely correlations with only few evidence of causality [62,63]. Concerning anxiety, we identified one bacterial genus, *Acidaminococcus*, with decreased abundance in BED patients with anxiety compared to BED patients without anxiety. We further show that the level of *Acidaminococcus* is negatively correlated to the HADS score in BED patients. Further experiments would now be require to delineate the potential causal role of this genus in the onset or maintenance of anxiety. Concerning depression, it is interesting to note that we observed in BED patients increased levels of *Eggerthella* and *Streptococcus*, that were also observed in patients with depressive disorder [64,65]. The *Eggerthella* genus, more particularly, has been reproducibly observed as enriched in many psychiatric disorders including MDD, bipolar disorder and schizophrenia [64]. Accordingly, we observed that patients from the “binge-eating” group with high levels of *Eggerthella* have a higher HADS Depression score. We also identified a positive correlation between *Eggerthella* levels and the HADS Depression score in our global cohort, as well as a negative correlation with the SF36 PCS score, reflecting a decrease in quality of life (Fig. 7). Additional taxa were identified as correlating with quality of life in AN or BED patients. These correlations were obtained after correction for BMI, suggesting that these associations are independent of the severity of BMI variations in these ED.

Cross-sectional studies of gut microbiota composition is a valuable approach to better understand the potential role of intestinal micro-organisms in a given disease. However, further work using longitudinal data are required to identify bacterial taxa correlating with patients’ clinical improvement or relapses and to define new prognostic markers in EDs. Longitudinal analyses would also be valuable to determine whether a gut dysbiosis precedes the onset of ED symptoms or is a secondary consequence of the changes in patients dietary of lifestyle habits. Such longitudinal studies have already been performed in the case of AN but are still lacking for other EDs [43,44,47,66–68]. Several studies have shown that weight normalization is not associated with a “normalization” of the gut microbiota composition [43,44,47,67,68]. This latent dysbiosis may contribute to the relapses and chronicization of AN in patients.

In complement to all these descriptive works, experimental approaches using animal models are urgently needed to demonstrate the causal role of gut microbiota changes in ED patients in the alteration of eating behaviour and in the onset or maintenance of ED-related comorbidities. Some preliminary studies have started tackling this issue. Gut microbiota alteration has been observed in rodent models of AN, which appears to be mostly starvation-related, and which is restored after refeeding [27,69,70]. Engraftment of germ-free mice with stool samples from AN patients have been used to address the causative role of patients’ microbiota on weight regulation and eating behaviour, albeit with mixed results [46,71,72]. In addition to these studies on AN, several studies addressed the role of the gut microbiota in the hedonic food intake of highly palatable food, which may contribute to binge eating or overeating, using various animal models [10,15,73–76]. These early studies now need to be completed and to include all types (and subtypes) of EDs to fully characterize the differential role of the gut microbiota in this set of diseases.

## Conclusion

Our work demonstrates that eating disorders are associated with a specific gut bacterial signature. We identified several bacterial taxa associated with specific ED subtypes or with ED-related comorbidities. Longitudinal studies will help decipher whether these bacterial signatures may be useful prognostic markers for EDs. The causal role of these taxa in the onset or maintenance of ED symptoms now need to be validated using *in vivo* approaches. These different approaches are essential to confirm the potential interest of developing innovative therapeutic treaments aiming at modulating the gut microbiota composition of ED patients, which will complement actual treatments and, hopefully, increase the quality of life and recovery of these patients.

## Declarations

### Ethics approval and consent to participate

The study design has been approved by the French national committee “Informatique et liberté” (N° CNIL: 1787487) and the Ethics Committee (N°CPP/CE 002/2014).

### Consent for publication

Not applicable

### Availability of data and material

The 16S rRNA gene raw sequences are available in the SRA database (Bioprojet: PRJNA1267983).

### Authors’ contributions

NA, MQ, SG, VF, HL, MC, PD and MPT were involved in the recruitment of ED patients and healthy volunteers, and the collection of clinical data and fecal samples. JB, MG and TD were involved in DNA extraction from fecal samples and sample analysis. LBB performed bioinformatics analyses of the 16S rRNA gene sequencing data and provided guidance regarding biostatistical analysis. DR performed bioinformatics analyses of the 16S rRNA gene sequencing data and was responsible for preparing the initial draft of the manuscript. All authors read and approved the final manuscript.

## Acknowledgements

We thank the staff of the CIC-CRB 1404 (CHU Rouen) for their assistance with human sample management and processing.

