## Supplemental Table S1 for "Characterisation of changes in the gut microbiota associated with eating disorders"

|  | **no ED**  (n=73) | **all ED**  (n=181) | no ED  vs all ED | **Anorexia**  (n=35) | **Bulimia**  (n=18) | **Binge-eating**  (n=128) |
| --- | --- | --- | --- | --- | --- | --- |
| **Women/Men** | 59/14 | 155/26 | *p* = 0.35 | 34/1 | 16/2 | 105/23 |
| **Age** (years) | 34.9 ± 10.8 | 38.0 ± 14.1 | *p* = 0.23 | 31.9 ± 13.1 | 34.0 ± 15.7 | 40.3 ± 13.7 |
| **BMI** (kg/m^2^) | 21.8 ± 1.7 | 31.5 ± 10.5 | ***p* < 0.0001** | 15.8 ± 1.4 | 23.1 ± 4.4 | 36.9 ± 6.8 |
| BMI class |  |  |  |  |  |  |
| <18.5 | 0 % | 20.5 % |  | 100 % | 11.1 % | 0 % |
| 18.5-24.9 | 100 % | 8.3 % |  | 0 % | 55.6 % | 3.9 % |
| 25-29.9 | 0 % | 7.7 % |  | 0 % | 27.8 % | 7.0 % |
| >29.9 | 0 % | 63.5 % |  | 0 % | 5.5 % | 89.1 % |

**Table S1: Clinical features of ED patients included in the study**
