## Supplemental Table S2 for "Characterisation of changes in the gut microbiota associated with eating disorders"

|  | **Anorexia** | **Bulimia** | **Binge-eating** | Anorexia  vs Bulimia | Anorexia  vs Binge-eating | Bulimia vs Binge-eating |
| --- | --- | --- | --- | --- | --- | --- |
| **EDI-2** |  |  |  |  |  |  |
| subscore DT | 8.6 ± 6.3 | 13.3 ± 5.7 | 10.3 ± 4.6 | ***p =* 0.016** | *p* = 0.50 | *p* = 0.09 |
| subscore B | 1.9 ± 3.7 | 9.3 ± 6.1 | 5.8 ± 5.8 | ***p* < 0.0001** | ***p* = 0.0001** | *p* = 0.21 |
| subscore BD | 11.7 ± 6.1 | 14.3 ± 7.9 | 20.5 ± 6.3 | *p* = 0.76 | ***p* < 0.0001** | ***p* = 0.005** |
| subscore I | 10.9 ± 7.5 | 8.6 ± 7.7 | 9.9 ± 7.2 | *p* = 0.86 | *p* > 0.99 | *p* > 0.99 |
| subscore P | 6.8 ± 4.3 | 8.2 ± 4.8 | 5.6 ± 4.1 | *p* > 0.99 | *p* = 0.48 | *p* = 0.10 |
| subscore ID | 6.4 ± 4.3 | 5.1 ± 4.6 | 5.1 ± 4.1 | *p* = 0.79 | *p* = 0.39 | *p* > 0.99 |
| subscore IA | 9.3 ± 6.4 | 11.5 ± 6.5 | 8.4 ± 6.4 | *p* = 0.94 | *p* > 0.99 | *p* = 0.23 |
| subscore MF | 6.4 ± 5.5 | 4.1 ± 3.3 | 4.8 ± 4.4 | *p* = 0.64 | *p* = 0.47 | *p* > 0.99 |
| subscore AS | 7.0 ± 3.9 | 7.5 ± 2.7 | 6.1 ± 3.5 | *p* > 0.99 | *p* = 0.90 | *p* = 0.22 |
| subscore IR | 5.0 ± 6.5 | 5.9 ± 5.3 | 4.8 ± 5.2 | *p* = 0.92 | *p* > 0.99 | *p* > 0.99 |
| subscore SI | 7.4 ± 4.3 | 7.0 ± 5.1 | 6.2 ± 4.4 | *p* > 0.99 | *p* = 0.39 | *p* > 0.99 |
| global score | 86.0 ± 40.2 | 98.8 ± 35.8 | 85.8 ± 34.8 | *p* > 0.99 | *p* > 0.99 | *p* = 0.84 |
| **BSQ** |  |  |  |  |  |  |
| subscore ABE | 19.8 ± 7.8 | 24.8 ± 10.8 | 30.5 ± 8.9 | *p* = 0.21 | ***p* < 0.0001** | *p* = 0.18 |
| subscore BDL | 30.1 ± 15.1 | 43.8 ± 15.1 | 46.6 ± 11.5 | ***p =* 0.015** | ***p* < 0.0001** | *p* > 0.99 |
| subscore ULV | 4.9 ± 2.2 | 6.9 ± 2.8 | 5.0 ± 2.0 | ***p =* 0.026** | *p* > 0.99 | ***p =* 0.021** |
| subscore CIW | 15.0 ± 7.3 | 22.2 ± 5.3 | 17.7 ± 4.7 | ***p* < 0.0001** | *p* = 0.19 | ***p =* 0.001** |
| global score | 89.9 ± 37.9 | 125.0 ± 36.7 | 125.3 ± 30.2 | ***p =* 0.013** | ***p* < 0.0001** | *p* > 0.99 |
| **QUAVIAM** |  |  |  |  |  |  |
| global score | 298.6 ± 115.6 | 275.7 ± 91.7 | 277.0 ± 97.9 | *p* > 0.99 | *p* = 0.6861 | *p* > 0.99 |

**Table S2: EDI-2, BSQ and QUAVIAM scores of ED patients included in the study**

EDI-2 subscores: DT, drive for thinness; B, bulimia; BD, body dissatisfaction; I, ineffectiveness; P, perfectionism; ID, interpersonal distrust; IA, interoceptive awareness; MF, maturity fears; AS, asceticism; IR, impulse regulation; SI, social insecurity. BSQ subscores: ABE, avoidance and social shame of body exposure; BDL, bodily dissatisfaction with the lower parts of the body; ULV, use of laxatives and vomiting to reduce body dissatisfaction; CIW, cognitions and inappropriate behaviours to control weight. Kruskal-Wallis test with Dunn’s correction.
