## Supplemental Figures S1 to S3 for "Characterisation of changes in the gut microbiota associated with eating disorders"

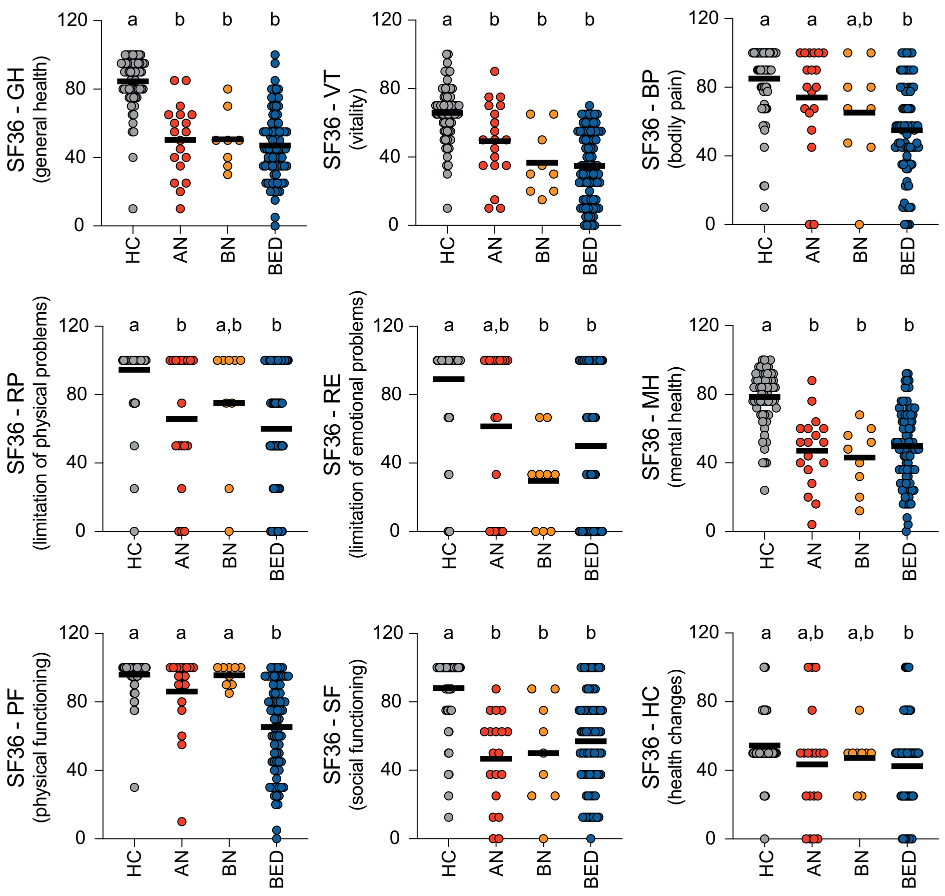


**Figure S1: SF36 subscores of ED patients and healthy controls.**
Individual values and means are represented (labeled means without a common letter differ; Kruskal-Wallis with Dunn's correction). HC, healthy controls; AN, «anorexia» group; BN, «bulimia» group; BED, «binge-eating» group.

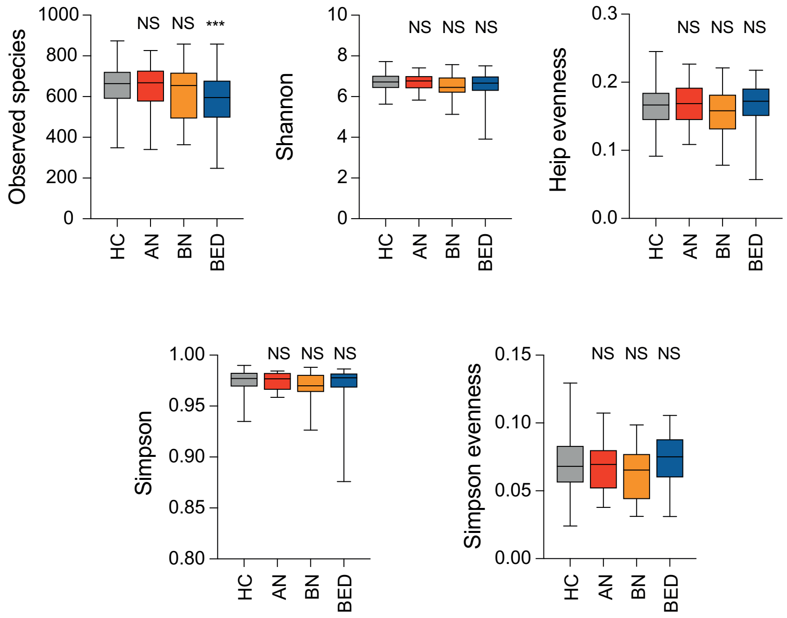


**Figure S2: Comparison of alpha diversity indexes between healthy individuals and ED patients**
The number of observed species, Shannon, Simpson, Heip evenness and Simpson evenness indexes in each group of individuals are represented as whisker plots with minimum and maximum values (NS, not signifiant; ***, *P*<0.001 vs healthy controls; Kruskal-Wallis test with Dunn's correction). HC, Healthy controls; AN, «anorexia» group; BN, «bulimia» group; BED, «binge-eating» group.

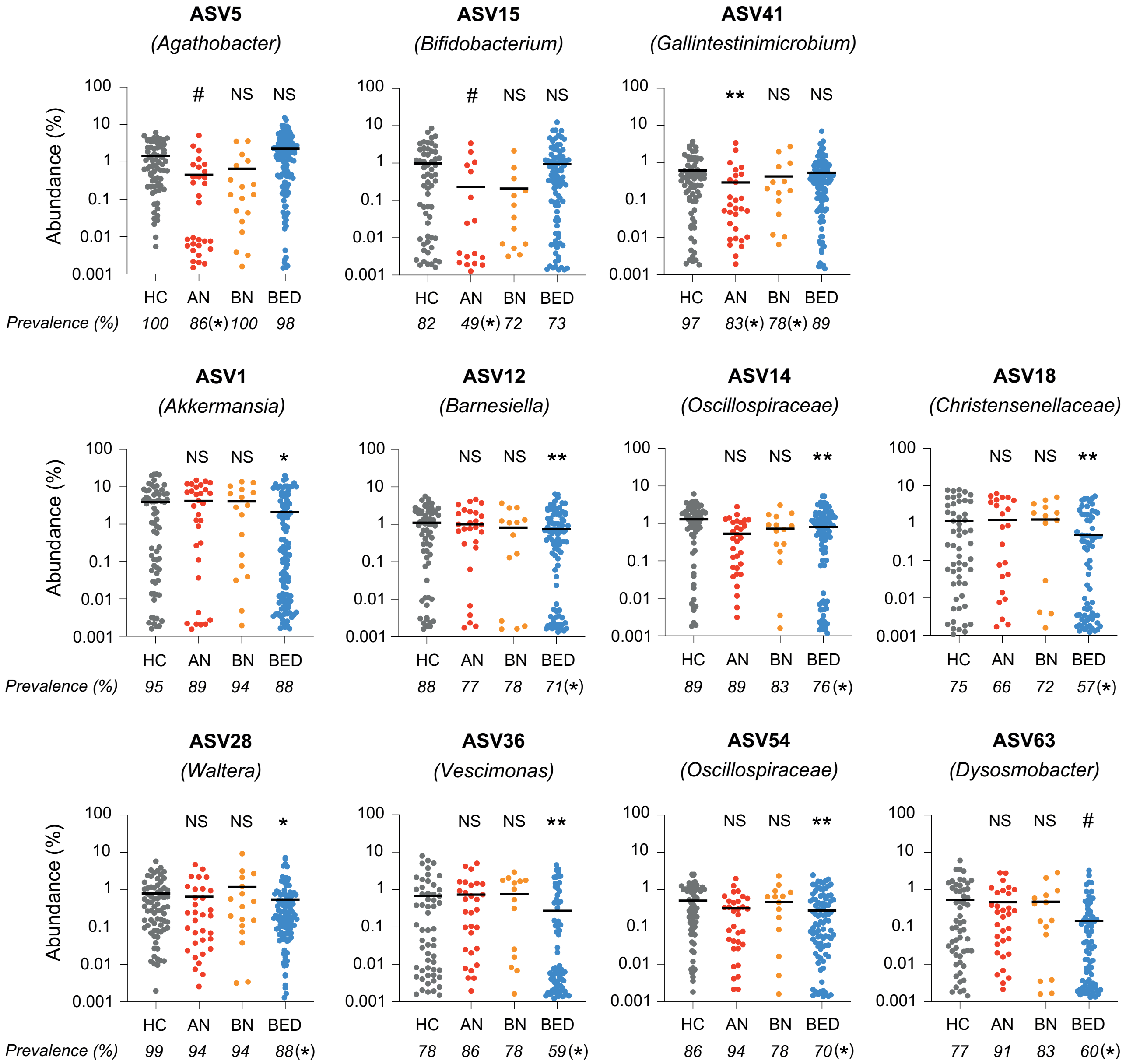


**Figure S3: Relative abundances of major ASVs altered in ED patients**
ASV with relative abundances above 0.5% in healthy individuals and showing significantly altered abundances in ED patients (*, $q<0.1$; **, $q<0.05$; Mann-Whitney; #, $q<0.1$ for both Mann-Whitney and Aldex2; comparison versus HC). HC, Healthy controls; AN, «anorexia» group; BN, «bulimia» group; BED, «binge-eating» group. The taxonomic identification of the ASV are indicated. Prevalences in each group are indicated below histograms (*, $p<0.05$; Fisher's exact test vs HC).
